# Bicarbonate Resensitization of Methicillin-Resistant *Staphylococcus aureus* to β-Lactam Antibiotics

**DOI:** 10.1101/570424

**Authors:** Selvi C. Ersoy, Wessam Abdelhady, Liang Li, Henry F. Chambers, Yan Q. Xiong, Arnold S. Bayer

**Affiliations:** Los Angeles Biomedical Research Institute, Torrance, CA 90502, USA; Geffen School of Medicine, University of California Los Angeles, Los Angeles, CA 90095, USA; Division of Infectious Diseases, Zuckerberg San Francisco General Department of Medicine, University of California, San Francisco School of Medicine, San Francisco, CA 94110 USA

**Keywords:** Methicillin-resistant *Staphylococcus aureus* (MRSA), Infective Endocarditis (IE) Antimicrobial Susceptibility Testing (AST), Beta-lactams (β-lactams), Sodium bicarbonate (NaHCO_3_), Penicillin-binding proteins (PBPs)

## Abstract

Endovascular infections caused by methicillin-resistant *Staphylococcus aureus* (**MRSA**) are a major healthcare concern, especially infective endocarditis (**IE**). Standard antimicrobial susceptibility testing (**AST**) defines most MRSA strains as ‘resistant’ to β-lactams, often leading to use of costly and/or toxic treatment regimens. In this investigation, five prototype MRSA strains, representing the range of genotypes in current clinical circulation, were studied. We identified two distinct MRSA phenotypes upon AST using standard media, with or without sodium bicarbonate (**NaHCO_3_**) supplementation: one highly susceptible to the anti-staphylococcal β-lactams, oxacillin and cefazolin (‘NaHCO_3_-responsive’) and one resistant to such agents (‘NaHCO_3_-nonresponsive’). These phenotypes accurately predicted clearance profiles of MRSA from target tissues in experimental MRSA IE treated with each β-lactam. Mechanistically, NaHCO_3_ reduced expression of two key genes involved in the MRSA phenotype, *mecA* and *sarA,* leading to decreased production of penicillin-binding protein (PBP) 2a (that mediates methicillin resistance), in NaHCO_3_-responsive (but not in NaHCO_3_-nonresponsive) strains. Moreover, both cefazolin and oxacillin synergistically killed NaHCO_3_-responsive strains in the presence of the host defense antimicrobial peptide (LL-37) in NaHCO_3_-supplemented media. These findings suggest that AST of MRSA strains in NaHCO_3_-containing media may potentially identify infections caused by NaHCO_3_-responsive strains that are appropriate for β-lactam therapy.

## INTRODUCTION

*Staphylococcus aureus* is a major bloodstream pathogen in both community-acquired and nosocomially-acquired scenarios, and is the leading cause of infective endocarditis (**IE**) in the industrialized world (1). Compounding the danger of S. *aureus* bloodstream infections (**BSIs**) is the steady rise of methicillin-resistant *Staphylococcus aureus* (**MRSA**) strains in many geographic regions in the US (2). MRSA is a serious infectious threat, causing more than 15,000 deaths in the U.S. each year (3).

MRSA have high minimal inhibitory concentrations (**MICs**) that are above Clinical Laboratory Standards Institute [CLSI] resistance breakpoints for most conventional β-lactam antibiotics, such as oxacillin, on standard antimicrobial susceptibility testing (**AST**) media. This finding implies a lack of efficacy of these agents in treating MRSA infections, as confirmed in selected experimental IE studies (4–6). Treatment options for MRSA are generally limited to costlier and/or more toxic drugs such as vancomycin, daptomycin, lipoglycopeptides, and fifth generation cephalosporins, such as ceftaroline (7–9). Additionally, great expense and effort have gone into development of such newer anti-MRSA drugs (10, 11).

AST protocols for MRSA have been standardized by the CLSI and involve growth of bacterial samples in 2% NaCl cation-supplemented, nutrient-rich Mueller-Hinton Broth (**CA-MHB**) (12, 13). However, MHB does not accurately represent the host milieu, and MICs observed in this medium are unlikely to mirror those exhibited by MRSA within specific host microenvironments. Although research efforts have been made to effectively model the host microenvironment *in vitro* or *ex vivo* (e.g., using simulated endocardial vegetations [SEVs]) (14, 15), these models are not suitable for large-scale AST in clinical laboratories.

Recently, several groups have attempted to improve standardized AST by altering growth conditions to better reflect the host environment. For example, AST of intracellular bacteria, such as *Salmonella,* in media that models the host phagolysosome can better predict treatment outcomes in murine bacteremia models (16, 17). Growth of bacteria in Dulbecco’s Modified Eagles Medium **(DMEM)** (but not in standard AST media) can stimulate expression of virulence factors typically exhibited *in vivo* (18, 19). These observations led to the discovery that AST of extracellular bacteria performed in such cell culture medium (including DMEM and RPMI) was a better predictor of *in vivo* treatment outcomes than standard AST media for *Acinetobacter* and some staphylococci (17, 20). Of note, Dorschner *et al.* identified the role of physiological concentrations of NaHCO_3_ in facilitating *in vitro* killing of *E. coli* and MRSA by host antimicrobial peptides (**AMPs**) (21). The carbonate molecule was capable of altering expression of a number of key regulatory genes in *E. coli*, including the homolog of the MRSA stress-response regulator, sigma factor B *(sigB).* Finally, our lab has recently confirmed the key role of NaHCO_3_ supplementation in rendering a range of bacterial pathogens, including MRSA, as more susceptible *in vitro* to multiple antibiotics, including β-lactams (17). The predictive role of this “NaHCO_3_-responsive” phenotype, *viz-a-viz* salutary outcomes to β-lactam therapy, was corroborated in a murine bacteremia model (17).

In the current study, we focused on the activity of two β-lactam agents routinely used for treating methicillin-susceptible S. *aureus* (**MSSA**) infections against five well-characterized prototype MRSA strains, in the presence or absence of NaHCO_3_-supplemented standard media. Using this modified AST schema, we demonstrate that NaHCO_3_ supplementation increased the *in vitro* susceptibility of selected MRSA strains to both cefazolin and oxacillin (i.e., ‘NaHCO_3_-responsive MRSA’). Although these drugs are typically not recommended for the treatment of MRSA (22), we observed that they can be highly effective in treating such NaHCO_3_-responsive MRSA strains in a rabbit model of IE. In contrast, MRSA strains that were NaHCO_3_-nonresponsive *in vitro* were recalcitrant to such β-lactam therapy *in vivo.*

If verified in larger MRSA screening studies, these novel findings may potentially prompt modifications of AST for MRSA; this, in turn, could potentially guide new treatment algorithms for selected MRSA infections with β-lactam agents such as cefazolin and anti-staphylococcal penicilins. Finally, mechanistically, NaHCO_3_-responsiveness correlated with the capacity of this molecule to suppress at least two key regulatory pathways intimately involved in the MRSA phenotype: *mecA*-PBP2a and *sarA* (5, 23, 24).

## RESULTS

### Effect of NaHCO_3_ on β-lactam susceptibility *in vitro*

MIC values obtained in standard CA-MHB for the five study strains were generally within 2-fold than those obtained in CA-MHB-Tris (Table S1). MRSA strains 11/11 (USA300 genotype) and MW2 (USA400 genotype) displayed a substantial decrease in MICs to cefazolin and oxacillin when grown in media containing 44 mM NaHCO_3_ (**Table 1**). Strain 11/11 also displayed a decrease in MICs to cefazolin and oxacillin, albeit less pronounced, when exposed to a more physiologically-relevant concentration of NaHCO_3_ (25 mM). In contrast, β-lactam MICs for three other prototype MRSA strains, COL (USA100 genotype), BMC1001 (USA500 genotype), and 300-111 (CC8, *spa* type 4, ‘Iberian clone’) were unaffected by exposure to NaHCO_3_ at either concentration (**Table 1**). Based on these two distinct phenotypes, we termed the two strains above, whose β-lactam MICs were substantially reduced by NaHCO_3_, as “responsive”, while the other three strains whose β-lactam MICs were unaffected by NaHCO_3_, as “nonresponsive”.

**Table 1.**
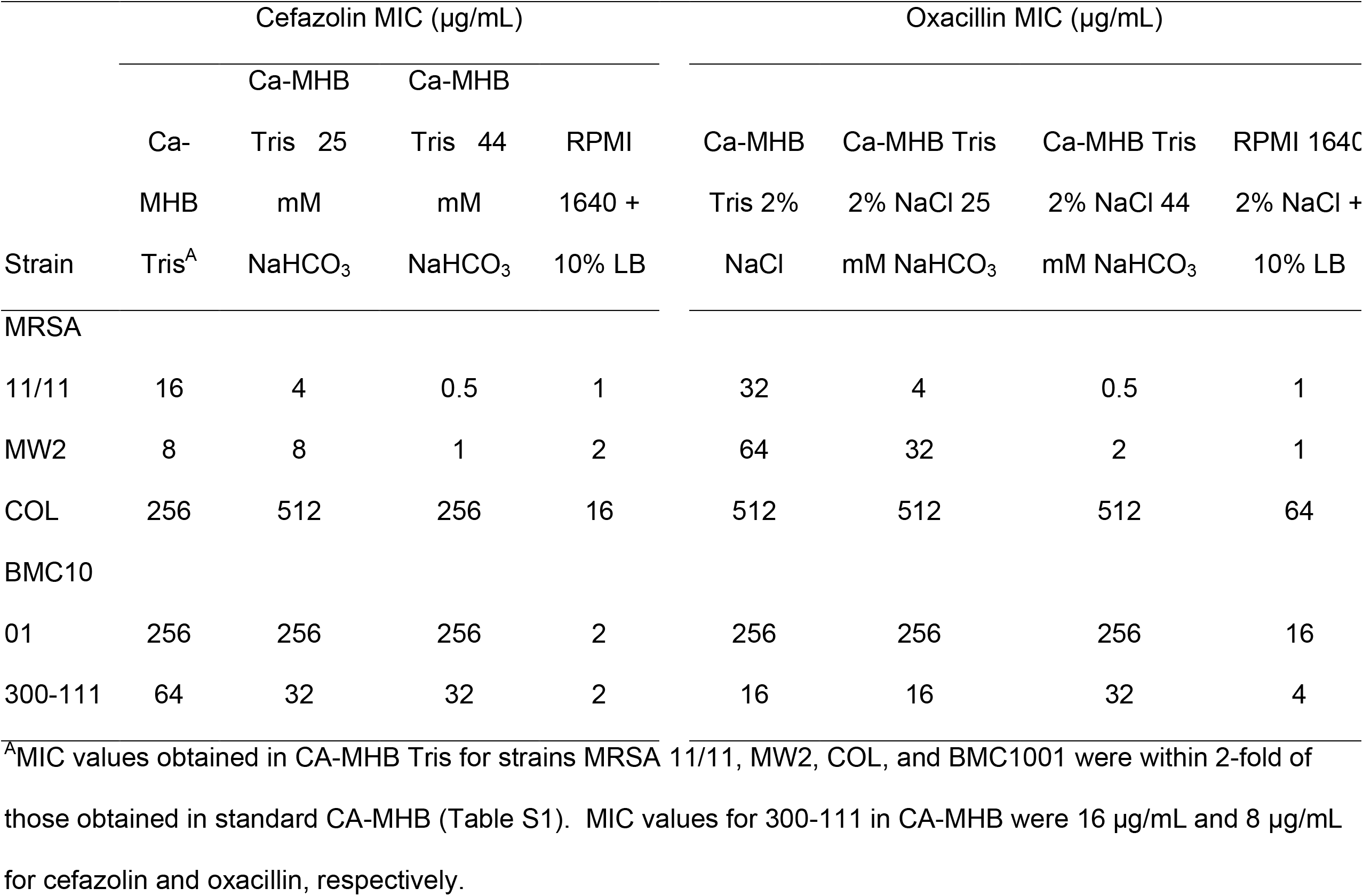
Minimum Inhibitory Concentration (MIC) of β-lactam antibiotics against methicillin-resistant *Staphylococcus aureus* grown in media with and without bicarbonate.

In addition, several recent investigations have promoted the use of the tissue culture medium, RPMI 1640, as a more physiologic, host-mimicking media and a better predictor of *in vivo* susceptibility to various antimicrobials than standard growth media, especially for Gram-negative bacteria (20, 25). As opposed to our NaHCO_3_ assays, we found that MICs to both β-lactams were decreased in RPMI for all five of our study strains, although the effect was somewhat less substantive for strain COL (**Table 1**).

To evaluate whether bicarbonate sensitization of MRSA to β-lactams was merely a ‘weak acid’ effect, we investigated the impact of salicylic acid exposure on β-lactam MICs among four of our study strains. Exposure of these MRSA to 25 and 50 μg/mL salicylic acid, concentrations that are physiologically achievable during aspirin therapy (26), had no effect on β-lactam MICs in any strain tested (**Table S1**). These data are inline with those of Farha *et al.* using two other weak acid molecules, acetate and borate (27). Taken together, this indicates that β-lactam re-sensitization of some MRSA in the presence of NaHCO_3_ is not the result of a generalized weak acid effect.

To further quantify the effect of NaHCO_3_ on MRSA β-lactam susceptibility profiles, a time-kill assay was conducted using log phase “responsive” MRSA 11/11 or “nonrespons¡ve” COL cells. As predicted by the MIC data, MRSA 11/11 displayed significantly greater killing when exposed to cefazolin and oxacillin in NaHCO_3_-containing media as compared to NaHCO_3_-free media (**Fig 1A**). After a 24 h incubation, a ≥ 4 log_10_ CFU/ml reduction in counts was observed for MRSA 11/11 exposed to 8 μg/mL cefazolin or 15 μg/mL oxacillin in NaHCO_3_-containing vs NaHCO_3_-free media. These latter β-lactam concentrations represent sublethal concentrations as determined in multiple pilot time-kill studies carried out in NaHCO_3_-free media. In contrast, strain COL displayed a minimal reduction in counts when grown in NaHCO_3_-containing vs. NaHCO_3_-free media under all testing conditions (**Fig 1B**). Similar differential time-kill results were obtained for the NaHCO_3_-responsive strain MW2 and the NaHCO_3_-nonresponsive strain, BMC1001 (**Fig S1**). Interestingly, the nonresponsive strain, 300-111, that has a relatively low baseline level of resistance to cefazolin and oxacillin, displayed an intermediate level of killing when exposed to 8 μg/mL cefazolin or 15 μg/mL oxacillin in NaHCO_3_-containing vs NaHCO_3_-free media (**Fig S1**).

**Figure 1.**
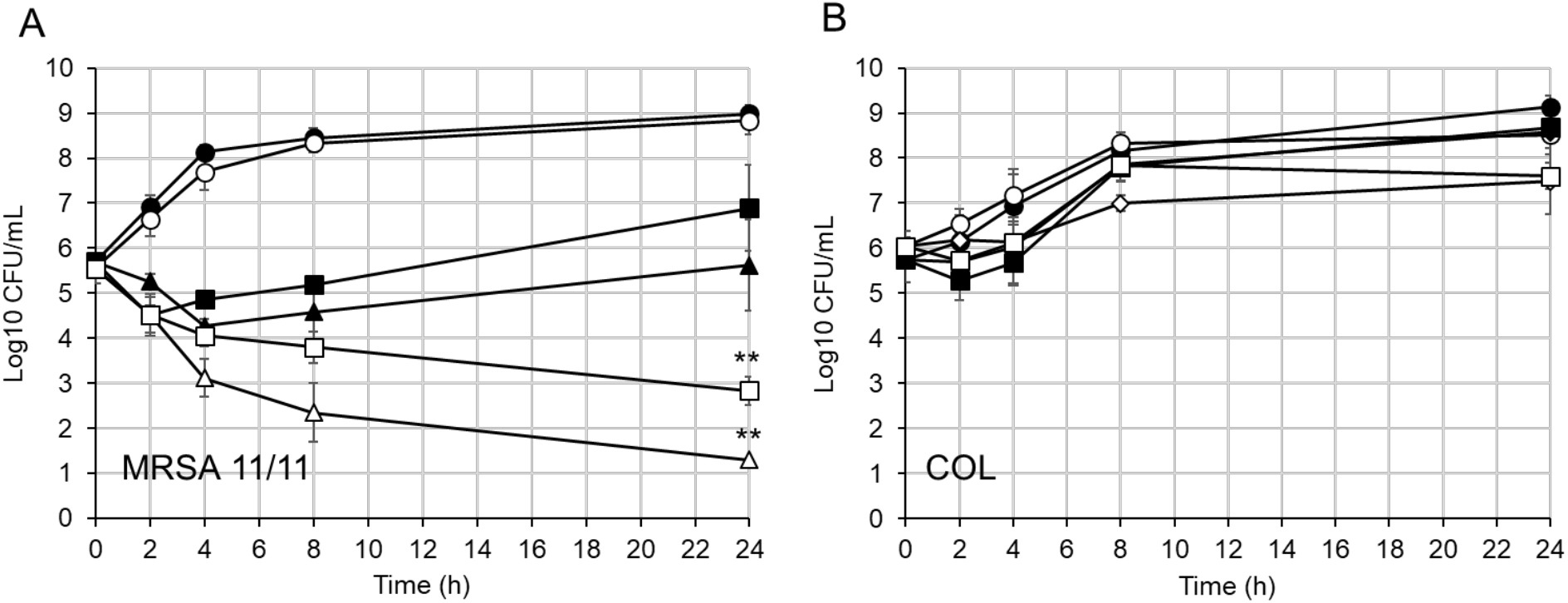
Time kill of log phase cells grown in media with and without bicarbonate. **(A)** MRSA 11/11 **(B)** COL. Growth in CA-MHB Tris (closed symbols), growth in CA-MHB Tris 44 mM NaHCO_3_ (open symbols), no drug (circles), 8 μg/mL cefazolin (triangles), 32 μg/mL cefazolin (diamonds), 15 μg/mL oxacillin (squares). The data are the means of two independent runs performed in triplicate for each condition ± the standard deviation. Statistical comparisons were made using a Kruskal-Wallis Single Factor ANOVA and post hoc Pairwise Mann-Whitney *U* test. Asterisks represent comparisons at 24 hour time point of MRSA 11/11 exposed to 8 μg/mL cefazolin in CA-MHB Tris vs CA-MHB Tris + 44 mM NaHCO_3_ and 15 μg/mL oxacillin in CA-MHB Tris 2% NaCl vs CA-MHB Tris 2% NaCl + 44 mM NaHCO_3_, **P < 0.01.

Supplementation with 44 mM NaHCO_3_ had only a modest effect on 24 hr growth kinetics for the five study strains. (**Fig S2**). With the exception of COL and 300-111, stationary phase cell counts at 24 h were not significantly different between media with and without 44 mM NaHCO_3_ for any strain tested.

### Population analyses in bicarbonate-containing media

Population analysis profiles (**PAPs**) are a standard quantitative *in vitro* assessment of the proportions of antibiotic-resistant sub-populations within a given strain vs specific antibiotics (28). MRSA strains contain a variable proportion of highly β-lactam-resistant subpopulations. For example, homogeneously-resistant (homo-resistant) strains usually contain a high percentage (e.g., > 10%) of such subpopulations, whereas heterogeneously-resistant (hetero-resistant) strains generally contain a lower percentage (e.g., < 0.01%) of resistant subpopulations.

To determine the effect of NaHCO_3_ on β-lactam-resistant subpopulations, PAPs for cefazolin were performed on our five prototype MRSA strains in NaHCO_3_-containing vs. NaHCO_3_-free agar. NaHCO_3_-responsive strains MRSA 11/11 and MW2 displayed hetero-resistant PAP phenotypes on NaHCO_3_-free agar, whereas NaHCO_3_-nonresponsive strains, COL, BMC1001, and 300-111 each displayed a more homo-resistant PAP phenotype (**Fig 2A**). To better characterize β-lactam hetero-resistant vs. homo-resistant phenotypes, using COL as our benchmark homo-resistant strain, we calculated the area under the PAP curve **(AUC)** ratios for MRSA 11/11, MW2, BMC1001, and 300-111 with respect to the COL AUC. AUC ratios of NaHCO_3_-responsive strains were significantly lower in the presence of NaHCO_3_ (compare **Fig 2A** to **Fig 2B**). Although NaHCO_3_ did have a slight repressive effect on the resistant sub-population of 300-111, the AUC for this strain in 44 mM NaHCO_3_ was significantly greater than both MRSA 11/11 and MW2. NaHCO_3_ had no effect on the AUC ratio of BMC1001 or the proportion of highly-resistant COL or BMC1001 cells (**Fig 2B**). Exposure to oxacillin, with or without NaHCO_3_, yielded similar results to cefazolin, although the magnitude of suppression of the highly-resistant subpopulations of the two NaHCO_3_-responsive strains (MRSA 11/11 and MW2) was less than that seen with cefazolin, and the differences were not statistically significant (data not shown).

**Figure 2.**
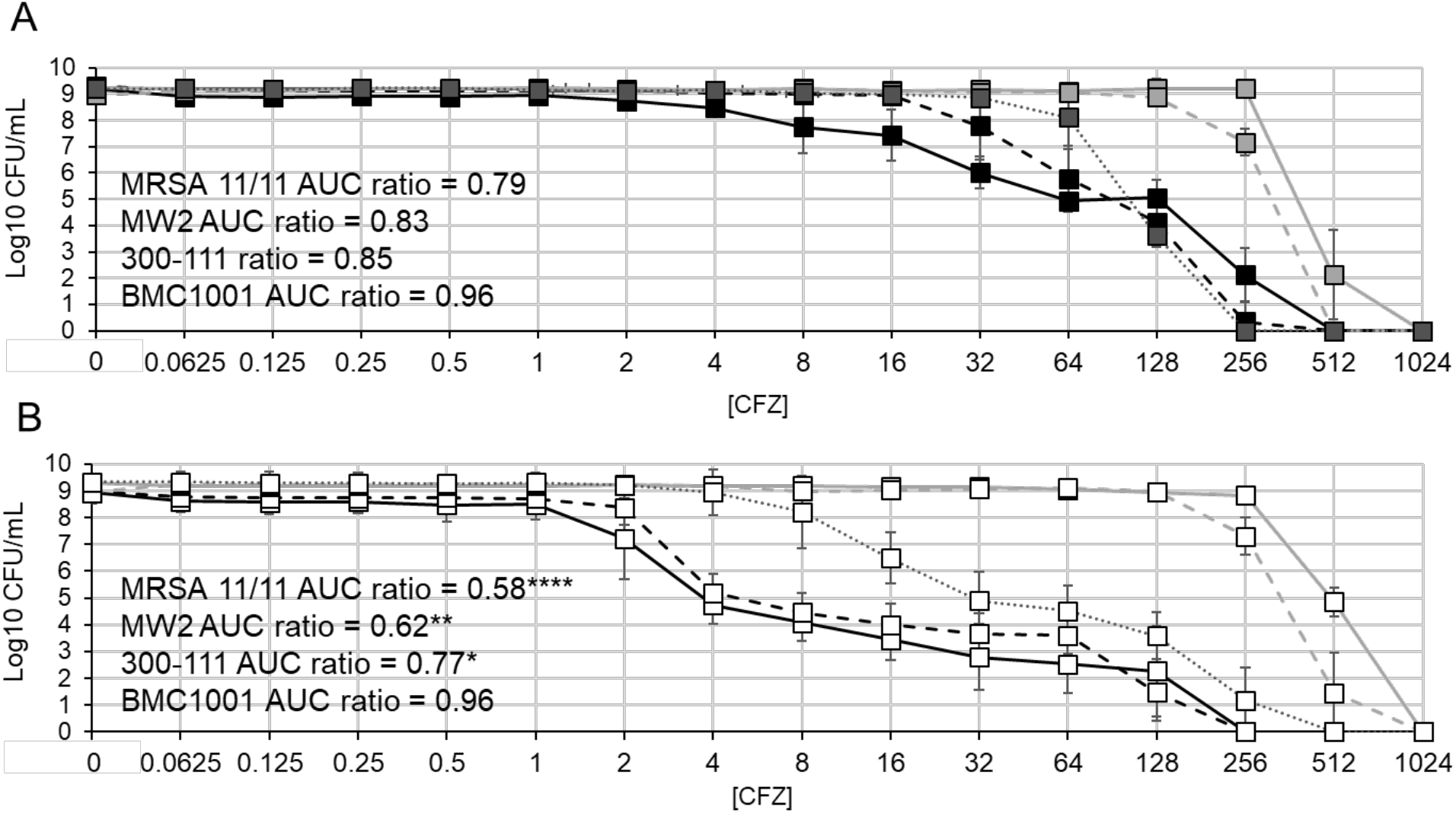
Population analysis of cells grown in media with and without bicarbonate. **(A)** Mueller-Hinton Agar (MHA) supplemented with Tris (closed symbols) **(B)** MHA supplemented with Tris and 44 mM NaHCO_3_ (open symbols). MRSA 11/11 (solid black line), MW2 (dashed black line), 300-111 (dotted grey line), COL (solid grey line), BMC1001 (dashed grey line). Drug abbreviation: cefazolin (CFZ). Area under the curve (AUC) ratios were compared to COL as a homo-resistant reference strain. The data are the means of two independent runs performed in triplicate for each condition ± the standard deviation. Statistical comparisons were made using a Kruskal-Wallis Single Factor ANOVA and post hoc Pairwise Mann-Whitney *U* test. MRSA 11/11, MW2, and 300-111 AUC in media containing 44 mM NaHCO_3_ is significantly reduced compared to media without NaHCO_3_, *P < 0.05, **P < 0.01, ****P < 0.0001; the AUC ratio of 300-111 in media containing 44 mM NaHCO_3_ is significantly greater than MRSA 11/11 and MW2, *P <0.05.

### Rabbit IE treatment outcomes with β-lactams

To verify the *in vivo* translatability of the NaHCO_3_ responsivity phenotypes determined *in vitro,* a rabbit model of aortic valve IE, treated with the two study β-lactams, was used. β-lactam treatment regimens were based on: i) published protocols for experimental IE therapy that are capable of clearing MSSA strains from target tissues (5, 29); ii) human-mimicking pharmacokinetic profiles (5, 29) (**Table S2**); and iii) our own pilot treatment outcome studies of experimental MSSA IE with these regimens (**Fig S4**). As predicted by *in vitro* MICs performed in NaHCO_3_-containing media, both NaHCO_3_-responsive strains were highly susceptible to β-lactam therapy *in vivo* to levels not dissimilar from that seen with the MSSA control strain ATCC 25923 (**Figs 3A; S4**). Thus, significant clearance of MRSA 11/11 and MW2 was observed in all target tissues sampled after 4 days of either cefazolin or oxacillin therapy. Of note, sterilization of multiple target tissues was seen in >70% of organ cultures from animals infected with the two NaHCO_3_-responsive when treated with cefazolin; in contrast, oxacillin therapy did not sterilize any target tissues despite significant reductions in MRSA counts in IE caused by these latter strains (data not shown). In contrast, β-lactam therapy was ineffective in reducing bacterial counts in the target tissues of rabbits infected with two NaHCO_3_-nonresponsive strains, COL or BMC1001 (**Fig 3B**). The *in vitro* NaHCO_3_-nonresponsive strain, 300-111, displayed a significant reduction in bacterial counts in all target tissues following cefazolin and oxacillin treatment; however the magnitutde of killing was significantly less than that observed in either NaHCO_3_-responsive strain (**Fig 3B**). Interestingly, the MICs determined in CA-MHB Tris containing 44 mM NaHCO_3_ were better predictors of treatment outcomes for all five strains than MICs determined in RPMI 1640, based roughly on previously established 2014 CLSI breakpoints [(cefazolin breakpoints are no longer employed in the current 2018 CLSI Guidelines) (13, 30): S ≤ 8 μg/mL, I = 16 μg/mL, R ≥ 32 μg/mL] and current oxacillin breakpoints: S ≤ 2 μg/mL, R > 4 ≥g/mL (31).

**Figure 3.**
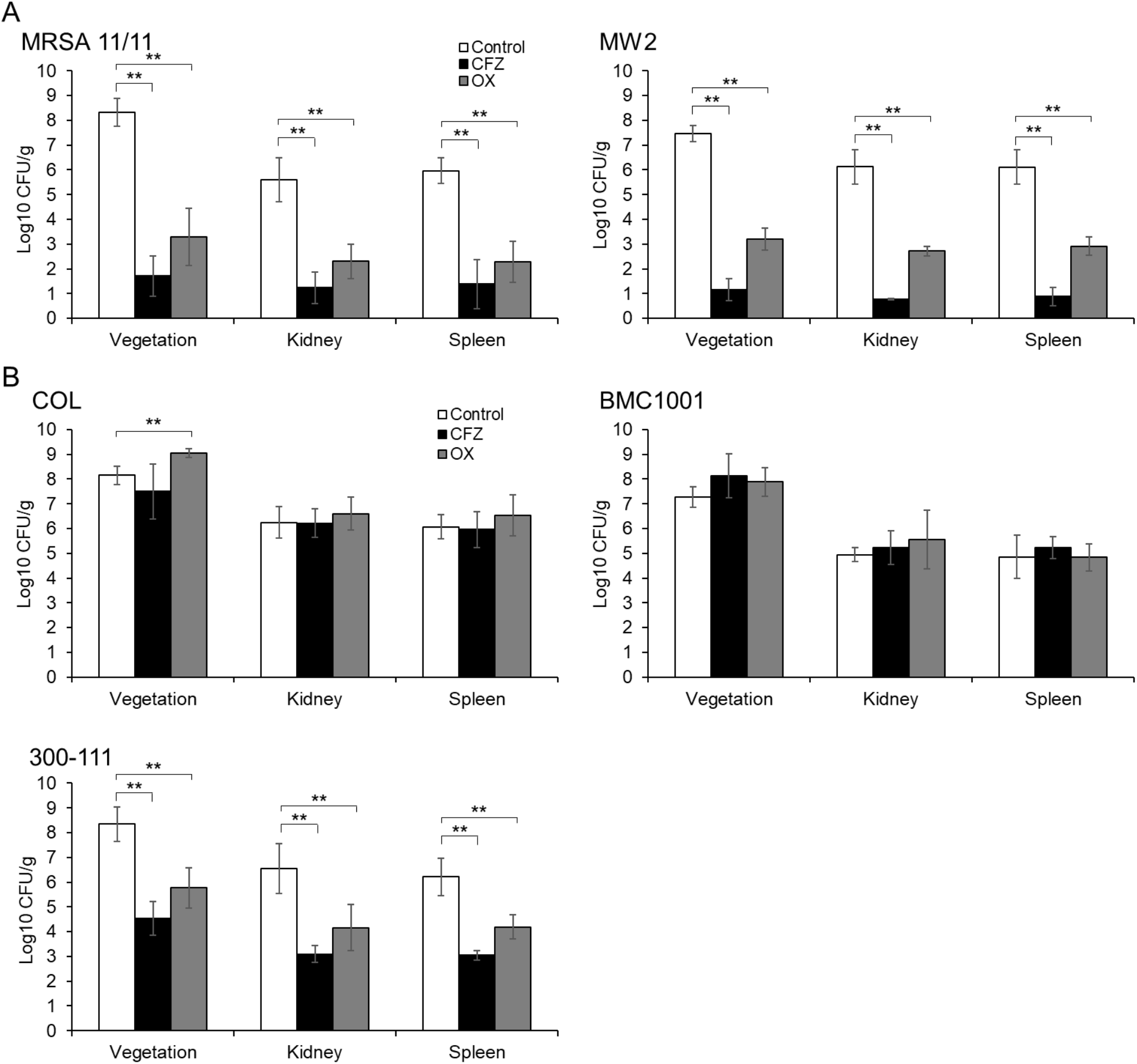
Treatment outcomes of rabbits with infective endocarditis treated with β-lactams. **(A)** NaHCO_3_-responsive strains. **(B)** NaHCO_3_-nonresponsive strains. All rabbits were treated with 100 mg/kg of cefazolin (CFZ) or oxacillin (OX), t.i.d., im, for 4 days. **MRSA 11/11**: control n = 6, CFZ n = 8, OX n = 6; **MW2**: control n = 7, CFZ n = 6, OX n = 7; COL: control n = 6, CFZ n = 7, OX n = 6; **BMC1001**: control n = 6, CFZ n = 7, OX n =7; **300-111**: control n = 10, CFZ n = 4, OX n = 6. The data presented are the mean tissue CFU/g for each treatment group ± the standard deviation. Statistical comparisons were made using a Kruskal-Wallis Single Factor ANOVA and post hoc Pairwise Mann-Whitney *U* test, **P < 0.01. 300-111 had significantly greater cell counts in all target tissues following cefazolin and oxacillin therapy compared to both MRSA 11/11 and MW2, *P < 0.05.

It was important to ensure that β-lactam therapy in experimental IE caused by the NaHCO_3_-responsive strains did not select for highly β-lactam-resistant subpopulations within target tissues post-β-lactam therapy. Thus, at the time of sacrifice, homogenates from these organs were parallel-plated onto TSA containing 64 μg/mL of either cefazolin or oxacillin in animals treated with these respective agents. No such high-level β-lactam-resistant colonies were detected for either MRSA 11/11-infected or MW2-infected animals following therapy with either of the two β-lactam agents (data not shown).

To investigate whether exposure to NaHCO_3_ itself might influence the tissue burdens of NaHCO_3_-responsive strains during induction of infection, in a separate study, MRSA 11/11 was grown overnight in CA-MHB Tris containing 25 mM or 44 mM NaHCO_3_ prior to infection of aortic-catheterized animals. At 24 h post-infection, there were no significant differences in target tissue MRSA counts between rabbits infected with MRSA 11/11 grown in NaHCO_3_-free CA-MHB Tris vs. 25 mM or 44 mM NaHCO_3_-containing CA-MHB Tris (**Fig S3B**). This confirmed that bicarbonate pre-exposure itself did not hinder the induction and early progression phases of infection in experimental IE caused by NaHCO_3_-responsive strains.

### NaHCO_3_ levels in experimental IE

To put our *in vivo* outcomes in experimental IE into perspective, we measured [HCO_3_^-^] levels in both infected and uninfected rabbits. Quantification of [HCO_3_^-^] concentrations in the blood of animals with IE vs. uninfected controls revealed that blood [HCO_3_^-^] levels remained relatively constant in the range of 20 – 25 mM (**Fig S3C**). Given that our maximal *in vitro* impact of NaHCO_3_ supplementation of standard media was seen at 44 mM vs 25 mM, this suggested that other factors are likely in-play *in vivo* which contribute to NaHCO_3_-responsivity in experimental IE (e.g., host immune molecules and/or cells).

### LL-37 synergy with β-lactams in the presence of physiological concentrations of NaHCO_3_

We hypothesized that the high-level killing exhibited by β-lactams *in vivo* against the two NaHCO_3_-responsive strains in the presence of physiologic concentrations of NaHCO_3_ (~20-25 mM) may be due to a synergistic effect between β-lactams and host immune factors, particularly host defense peptides. To further investigate this, we performed a cell survival assay utilizing sub-lethal concentrations of the human cathelicidin, LL-37, in combination with cefazolin or oxacillin, in a minimal medium containing 25 mM NaHCO_3_. Strains were exposed to LL-37 and the β-lactams alone or in combination, in the presence of this physiological concentration of NaHCO_3_.

When exposed to a combination of LL-37 and either cefazolin or oxacillin, both NaHCO_3_-responsive strains displayed significantly lower MRSA survivals as compared to either β-lactam agent alone (**Fig 4A-B**). In contrast, the NaHCO_3_-nonresponsive strain, COL, did not display synergistic killing when exposed to a combination of LL-37 plus either cefazolin or oxacillin (**Fig 4C**), consistent with results in the IE model. These data indicate that, although higher concentrations of NaHCO_3_ (44 mM) are required *in vitro* to disclose “responsiveness” among MRSA strains, host defense and/or other serum factors may foster such responsivity *in vivo* in combination with more physiologic NaHCO_3_ levels (~20-25 mM).

**Figure 4.**
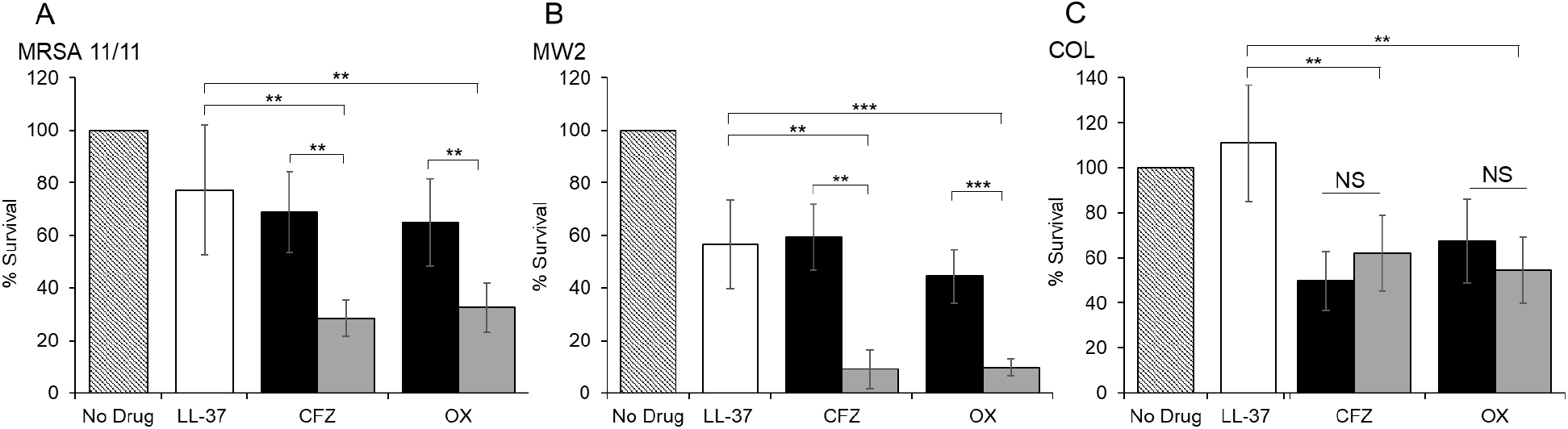
LL-37 synergy with B-lactams in media containing 25 mM NaHCO_3_. **(A)** MRSA 11/11 (NaHCO_3_-responder). **(B)** MW2 (NaHCO_3_-responder). (C) COL (NaHCO_3_-nonresponder). Drug abbreviations: cefazolin (CFZ), oxacillin (OX). No Drug (hatched); LL-37 only (white); CFZ or OX (black); CFZ or OX + LL-37 (grey). MRSA 11/11 was exposed to 2.5 μg/mL LL-37 and 0.03125 μg/mL CFZ or OX. MW2 and COL were exposed to 5 μg/mL LL-37 and 0.0625 μg/mL CFZ or OX. Percent survival calculated after 4 h exposure to antimicrobials in bicarbonate-free DMEM, supplemented with 25 mM NaHCO_3_. The data are the means of three independent runs performed in triplicate for each condition ± the standard deviation. Statistical comparisons were made using a Kruskal-Wallis Single Factor ANOVA and post hoc Pairwise Mann-Whitney *U* test, **P < 0.01, ***P < 0.001.

### Mechanisms of NaHCO_3_-mediated β-lactam resensitization of MRSA

#### *mecA* and *sarA* expression

To understand the potential genetic basis for altered β-lactam susceptibility in selected MRSA strains in NaHCO_3_-containing media, we investigated the influence of NaHCO_3_ on *mecA* and *sarA* gene expression. The *mecA* locus encodes penicillin-binding protein 2a (PBP2a) that confers β-lactam resistance in MRSA strains (24). In addition, recent studies have demonstrated that the global virulence gene regulator, *sarA,* can also modulate β-lactam resistance via both *mecA-* dependent and *mecA*-independent mechanisms (23). To investigate the influence of NaHCO_3_ exposure on *mecA* and *sarA* gene expression, RNA was extracted from cells grown in media with and without NaHCO_3_ in the absence or presence of ½ MIC of oxacillin (to maximally induce *mecA* expression). The qRT-PCR analyses revealed that *mecA* and *sarA* gene expression were each significantly repressed in the two NaHCO_3_-responsive strains under both these *mecA*-noninducing and -inducing conditions (**Fig 5A;B**). In contrast, *mecA* gene expression was not repressible in any NaHCO_3_-nonresponsive strain (**Fig 5A;B**). The expression of *sarA* was only slightly repressed in COL in NaHCO_3_-supplemented media with oxacillin induction, while being non-repressible in media without oxacillin induction (**Fig 5A;B**). The expression of *sarA* was also not repressible in the other NaHCO_3_-nonresponsive strains, BMC1001 and 300-111 (**Fig 5A;B**).

**Figure 5.**
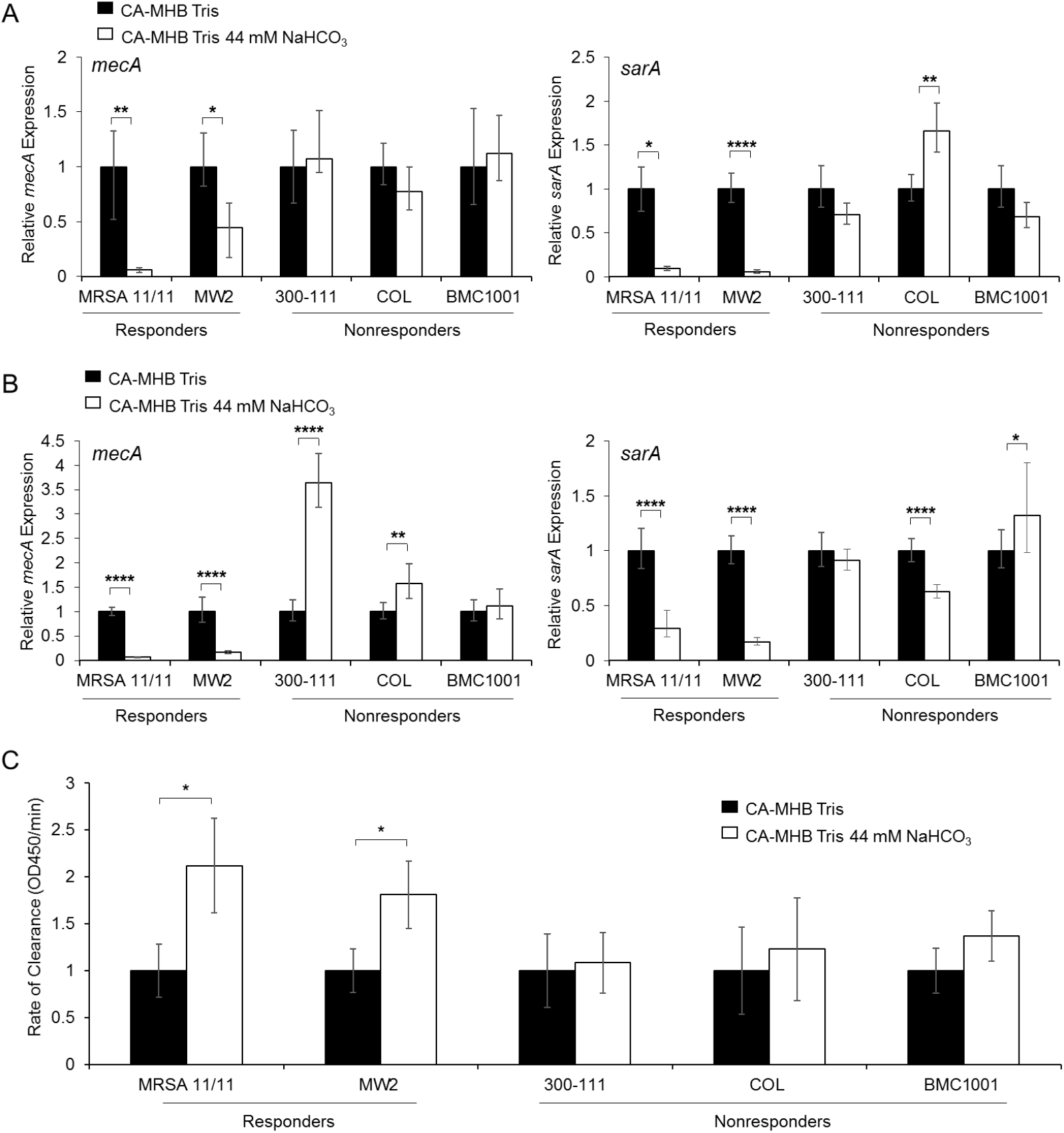
*mecA* and *sarA* expression in NaHCO_3_ responsive and non-responsive strains. **(A)** qRT-PCR of *mecA* and *sarA* gene expression in stationary phase cells grown in CA-MHB Tris with and without NaHCO_3_. **(B)** qRT-PCR of *mecA* and *sarA* gene expression in stationary phase cells grown in CA-MHB Tris 2% NaCl with and without NaHCO_3_ with ½ MIC concentration of oxacillin. Gene expression levels are normalized to the housekeeping gene *gyrB.* Gene expression values normalized to CA-MHB Tris for each strain. All data are derived from two independent biological replicates, tested in triplicate on two separate occasions. **(C)** Lipase production analysis in NaHCO_3_-responsive, nonresponsive, and *ΔsarA* mutant strains grown in media with and without NaHCO_3_. Values normalized to CA-MHB Tris for each strain. The data are the means of two independent runs performed in triplicate for each condition ± the standard deviation. Statistical comparisons were made using a Kruskal-Wallis Single Factor ANOVA and post hoc Pairwise Mann-Whitney *U* test, *P < 0.05, **P < 0.01, ****P < 0.0001.

To phenotypically verify that the *mecA* repression observed in NaHCO_3_-responsive strains correlated with diminished PBP2a protein production, a macro-agglutination PBP2a production assay was utilized. A clear reduction in PBP2a agglutination was observed in both NaHCO_3_-responsive strains after growth in NaHCO_3_-containing media (**Table 2, Fig S3**), consistent with the reduced expression of *mecA.* The NaHCO_3_-nonresponsive strains, COL and BMC1001, displayed high levels of agglutination in media with and without NaHCO_3_ exposures (**Table 2, Fig S3**). Although *mecA* gene expression was not repressible in strain 300-111 in the presence NaHCO_3_, the level of observable PBP2a agglutination was slightly diminished in media containing NaHCO_3_.

**Table 2.**
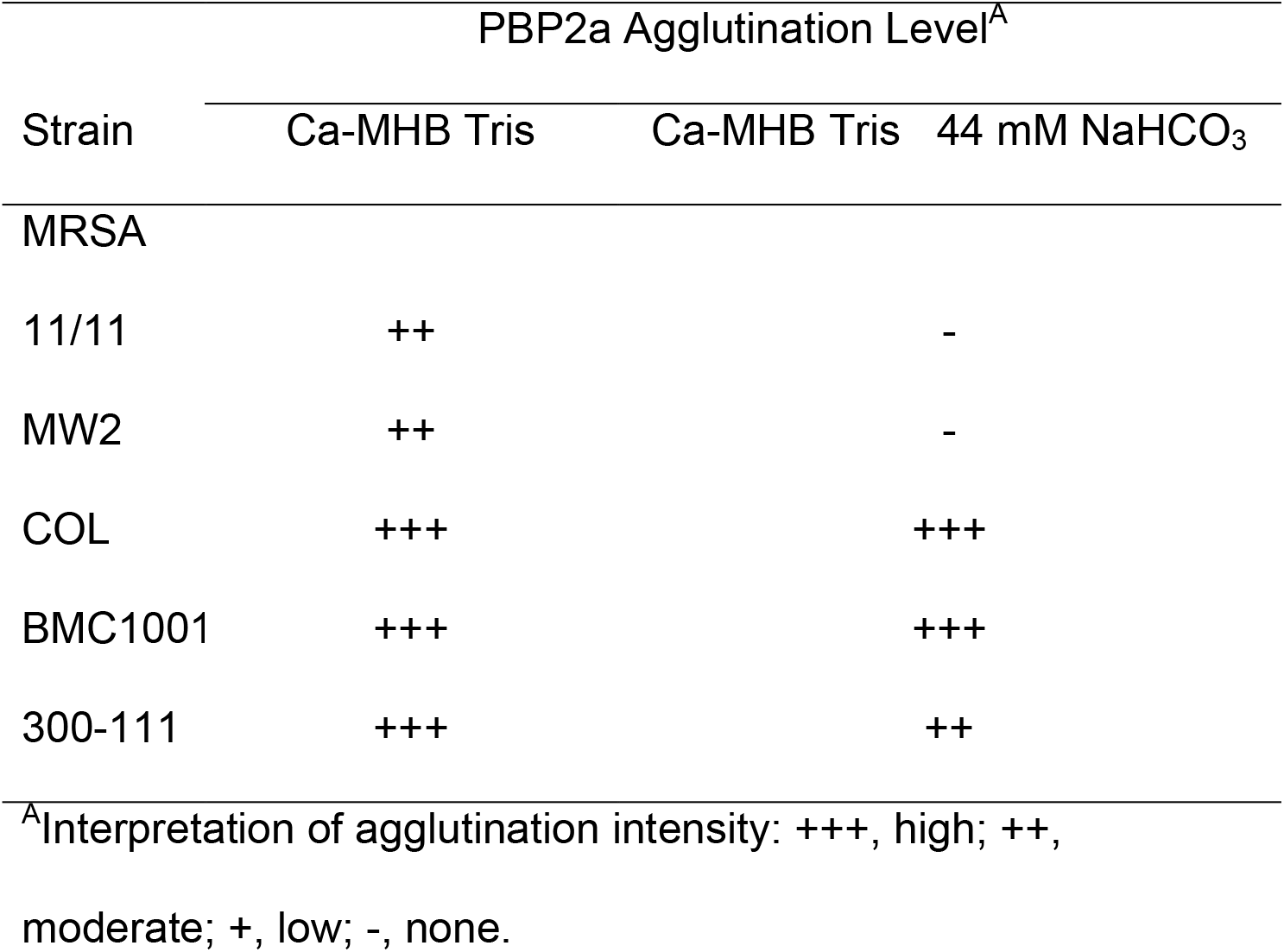
PBP2a agglutination of NaHCO_3_-responsive and nonresponsive strains grown in media ± 44 mM NaHCO_3_.

Lipase production is one of the signature phenotypes normally repressed by the global regulator *sarA* (32, 33). To phenotypically confirm the blunting of *sarA* gene expression by NaHCO_3_ exposures seen in NaHCO_3_-responsive vs. -nonresponsive strains, a tributyrin clearance assay was used to measure lipase production. As predicted by NaHCO_3_ repression of *sarA* gene expression, NaHCO_3_-responsive strains had significantly higher levels of lipase activity when exposed to NaHCO_3_ (**Fig 5C**). In contrast, lipase production in NaHCO_3_-nonresponsive strains was not affected by the presence of NaHCO_3_ (**Fig 5C**).

## DISCUSSION

Several studies have demonstrated the importance of NaHCO_3_, the body’s primary biological buffer, in altering susceptibility of *S. aureus* to various antibiotics, including β-lactams, as well as to the prototypical host defense peptide, LL-37 (17, 21, 27). Herein, we show the ability of NaHCO_3_ to increase susceptibility in selected MRSA strains to two conventional β-lactam agents commonly used to treat MSSA infections. This raises the intriguing possibility of treating MRSA infections with such β-lactam agents, a concept not currently endorsed by any therapeutic guidelines for MRSA (22). Since such β-lactams are relatively inexpensive and exhibit infrequent serious side effects (34), our findings are of potential clinical relevance. The results show that certain MRSA strains found to be β-lactam-susceptible *in vitro* by AST with NaHCO_3_ supplementation, can be effectively treated by β-lactam therapy in a prototypical model of endovascular infections, experimental IE.

*In vitro,* NaHCO_3_-responsive MRSA strains have a typical signature. They are hetero-resistant on PAPs and have relatively low MICs within the MRSA range, to conventional β-lactams (i.e., to cefazolin and oxacillin). Phenotypically, as compared to NaHCO_3_-nonresponsive MRSA strains, such NaHCO_3_-responsive MRSA strains are killed significantly better *in vitro* by these β-lactams in the presence vs. absence of NaHCO_3_, and NaHCO_3_ substantially represses the growth of their resistant sub-populations.

Importantly, not all strains with relatively low intrinsic β-lactam MICs display the NaHCO_3_-responsive phenotype, as demonstrated by strain 300-111. Despite displaying nearly identical basal β-lactam MICs to MRSA 11/11 in standard media, this NaHCO_3_-nonresponsive strain was incompletely cleared from target tissues in experimental IE following cefazolin and oxacillin treatment. These outcomes underscore the concept that the resistant subpopulations of such low-MIC, but NaHCO_3_-nonresponsive, strains can not be simply eliminated by achieving supra-MIC β-lactam serum levels in such infections.

In our investigations, AST was performed at both 44 mM NaHCO_3_, a concentration present in certain cell culture medium (e.g., DMEM), as well as at 25 mM NaHCO_3_, a physiologically-relevant concentration for humans and rabbits (35). Although we saw highly effective β-lactam-mediated killing of our ‘responsive’ strains *in vivo*, we saw only a modest impact at physiologically-relevant concentrations of 25 mM NaHCO_3_ *in vitro.* These results suggested that other factors previously implicated in endovascular pathogenesis, such as host defense peptides [e.g. LL-37 or α-defensin, hNP-1; (36–38)], might synergize with β-lactams at physiological NaHCO_3_ concentrations to yield *in vivo* killing mirroring those observed *in vitro* at 44 mM NaHCO_3_. Of note, data, at least for LL-37, supported this notion. We recognize that other innate host immune factors (e.g., neutrophils; serum complement; other host peptides; anti-staphylococcal antibodies; etc.) may also contribute to enhanced bacterial killing by β-lactams *in vivo* in the presence of physiological concentrations of NaHCO_3_.

In terms of how physiological concentrations of NaHCO_3_ might impact MRSA to alter its intrinsic β-lactam susceptibility profiles, Farah *et al.* (29) proposed that this molecule was simply acting through its capacity to collapse the proton motive force (PMF = Δψ + ΔpH). Thus, [HCO_3_^-^], via Le Chatelier’s Principle, would drive the reaction, H+ + [HCO_3_^-^] >>>> H_2_0 + CO_2_ to consume protons, and collapse PMF by mitigating the ΔpH. In their study, they found that an MSSA strain became more resistant to oxacillin in the presence of NaHCO_3_ (27), and attributed their observations to globally decreased cellular respiration and growth rates. Since β-lactams are more effective against rapidly dividing cells, the authors postulated the mechanism to be based in an overall impact of reduced respiratory energy production to fuel growth in the presence of PMF-modifying concentrations of NaHCO_3_. In contradistinction, our study observed essentially the opposite result in MRSA; i.e., that certain MRSA strains become highly susceptible to β-lactams in the presence of NaHCO_3_, indicating that additional changes to MRSA-defining gene expression patterns may overcome any inhibitory effect of NaHCO_3_ on global cell metabolism to render such strains as β-lactam-susceptible.

The β-lactam treatment of rabbits infected with *in vitro* NaHCO_3_-responsive strains was highly effective in clearing infection in all target organs assessed. It was hypothesized that a decrease in overall virulence stimulated by growth in a NaHCO_3_-containing microenvironment might be part of the explanation for increased susceptibility to β-lactam antibiotics under these conditions. However, overnight growth of the NaHCO_3_-responsive strain, MRSA 11/11 in the presence of 25 or 44 mM NaHCO_3_ prior to infection had no effect on the strain’s ability to induce or propagate experimental IE.

Intriguingly, our *in vitro* MIC data obtained in CA-MHB containing 44 mM NaHCO_3_ was a better predictor of *in vivo* outcomes than the tissue culture medium, RMPI 1640. Although use of this latter medium for AST is gaining some traction as a “host mimicking milieu” [especially for Gram-negative pathogens (20, 25, 39)], we found that it falsely predicted efficacy of oxacillin and cefazolin for the *in vivo* treatment of some study MRSA strains, (e.g., BMC1001). Further head-to-head screening in both media, using larger MRSA strain collections, will be required to fully identify which medium is the best predictor of β-lactam efficacy *in vivo* against MRSA.

Gene expression analyses revealed that *mecA* and *sarA* were highly repressed *in vitro* in NaHCO_3_-responsive vs. NaHCO_3_-nonresponsive strains. Repression of the *mecA* gene was, in turn, found to directly correspond to a decrease in PBP2a protein expression. It is unclear, however, if NaHCO_3_ has a direct impact on *mecA* expression, or if altered *mecA* gene expression is mediated through repression of *sarA* or other global regulons. Recent studies have shown that deletion of *sarA* and *sigB* can reduce *mecA* expression (23); therefore NaHCO_3_ may be altering *mecA* expression directly and/or through one or both of these global regulatory pathways. In addition, we observed the *mecA* and *sarA* expression levels were reduced by NaHCO_3_ exposures, in both the presence or absence of oxacillin induction (**Figs 5A;B**). This raises an intriguing dual-mechanim notion that NaHCO_3_ can directly effect the basal expression of these genes, as well as interfere with *mecA* induction by β-lactams, such as oxacillin.

As discussed above, Dorschner *et al.* determined that NaHCO_3_ had a significant repressive impact on expression of the *E. coli* homologue of the key S. *aureus* stress-response regulon, *sigB* (21). This locus is known to regulate pigment production in S. *aureus* (40); thus, factors that repress *sigB* expression reduce carotenoid expression, altering membrane fluidity (41). Carotenoid is critical for oligomerization and proper insertion of PBP2a into the cell membrane (42). NaHCO_3_’s potentially repressive impact on the *sigB* regulatory axis, could thus result in a decrease in carotenoid production. This event, combined with decreased PBP2a expression in NaHCO_3_-responsive strains, could yield MRSA cells that are phenotypically “mecA-defective”. In this regard, pilot studies in our laboratory have confirmed that *in vitro* NaHCO_3_ exposures in ‘responsive’ MRSA can repress both *sigB* expression, as well as carotenoid production (data not shown). Additional experiments are required to identify the comparative influence of NaHCO_3_ on both *sigB* expression and carotenoid production in NaHCO_3_-responsive vs. NaHCO_3_-nonresponsive strains.

Although we have identified at least two genetic targets of NaHCO_3_ that may influence MRSA resistance to β-lactams, it is highly likely that NaHCO_3_ has pleotropic effects on gene expression which may contribute to β-lactam susceptibility in NaHCO_3_-responsive strains. Furthermore, different pathways may be activated or repressed in individual “responsive” and “nonresponsive” strains, resulting in multiple “genetic types” of NaHCO_3_-responsiveness. For example, wall techoic acid (WTA) forms a scaffold for PBP2a maturation, allowing its insertion into the cell membrane (43). Deletion of genes involved in WTA synthesis can render MRSA strains more susceptible to β-lactams (43, 44), indicating that this may be another potential site of NaHCO_3_ action on gene expression. The modest decrease in PBP2a expression in strain 300-111, despite *mecA* expression being non-repressible by NaHCO_3_ in this strain, highlights this latter point. Thus, NaHCO_3_ may be affecting multiple pathways in this nonresponsive strain, causing an increase in *mecA* gene expression, but a slight overall decrease in membrane insertion of mature PBP2a.

Another potential ‘checkpoint’ for NaHCO_3_ is PBP4, a protein involved in the generation of highly cross-linked peptidoglycan. Although this PBP is normally dispensable in MSSA (45, 46), MRSA strains require its activity for proper peptidoglycan cross-linking (47), without which it is dependent on the β-lactam-susceptible, PBP2, for peptidoglycan synthesis. Interestingly, inhibition of PBP4 activity has been shown to diminish cell wall cross-linking in β-lactam hetero-resistant MRSA strains, but had no effect on cross-linking in homo-resistant strains (48). Currently, little is known about the regulation of PBP4 expression (49); however, further investigations into the effect of NaHCO_3_ on PBP4 protein production may offer additional insights into mechanisms underlying NaHCO_3_-responsive vs NaHCO_3_-nonresponsive phenotypes.

In MRSA, there is a phenomenon somewhat akin to NaHCO_3_-β-lactam responsivity called the “see-saw” effect, in which MRSA cells that evolve daptomycin resistance become resensitized to β-lactams (50–53). This phenomenon involves the complex interaction between at least four genes. *First*, *prsA* is involved in the proper functioning of PBP2a by encoding a membrane-anchored chaperone required for proper PBP2a folding and insertion into the membrane (50, 51). *Second,* expression of *prsA,* in turn, is regulated by the two-component regulatory system, VraSR (52); in the see-saw effect, expression of *vraSR* is upregulated (50, 53). *Third, prsA* expression is also co-regulated by gain-in-function mutations in the *mprF* operon which is involved in cell membrane phospholipid synthesis (54). Of interest, in the see-saw effect, alteration of the cell membrane apparently prevents membrane insertion of PrsA, inhibiting this chaperone’s net functionality (50, 55). *Fourth,* expression of *mprF* is regulated by the sense-response two-component regulator, *graRS* (56), and is upregulated in daptomycin-resistant strains compared to daptomycin-susceptible strains (54, 57). Collectively, these data above provide a number of additional mechanistic sites potentially influenced by NaHCO_3_. Many of these genetic pertubation possibilities are under active investigation in our labs by whole genome sequencing and RNA sequencing analyses.

This work highlights the exciting possibility that current standard-of-care β-lactams that are low-cost and low-toxicity may be effective in treating infections caused by NaHCO_3_-responsive MRSA strains. Further work is needed to fully elucidate the mechanism of NaHCO_3_-induced β-lactam susceptibility, as well as to identify phenotypic and genetic “signatures” for NaHCO_3_-responsive MRSA strains. One of the major limitations of our study is the fact that only five MRSA strains were investigated. We are currently screening a large collection of well-characterized clinical MRSA strains which represent the broad range of clonal complex, *agr, SCCmec* and *spa* types in current world-wide circulation, for their NaHCO_3_-responsive profiles *in vitro.* Subsets of these strains will then be subjected to the same *in vivo* testing in the IE model as in the current work, to further verify the linkage between *in vitro* NaHCO_3_-responsivity and effective β-lactam therapy *in vivo.*

A key *in vivo* finding in the current study was that, in NaHCO_3_-responsive MRSA IE, neither treatment with oxacillin nor cefazolin selected for emergence of high-level (MICs > 64 μg/ml) β-lactam-resistant subpopulations within cardiac vegetations. It should be emphasized, however, that one additional limitation of the present investigation was that rabbit vegetations are considerably smaller than those of humans with IE (~2-3 mm in diameter, corresponding to a weight of ~50-100 mg vs. ~1 cm in diameter corresponding to ~500-1,000 mg, respectively) (58–60). Therefore, if bacterial densities observed in the rabbit IE model (which can reach 10^8^-10^9^ CFU/gm) are similar to those in humans, then the total bacterial burden in human vegetations would be at least 10 times higher than in the rabbit IE model. This would correspondingly increase the chance that highly-resistant organisms might well emerge during β-lactam therapy in human IE. This metric will need to be carefully monitored in any future clinical trials.Once a reliable and facile method for identifying potentially β-lactam-responsive MRSA strains has been established and verified, large-scale clinical trials to evaluate the effectiveness of β-lactam therapy for treating MRSA infections in human patients would be warranted.

## METHODS

### Bacterial strains and media

Methicillin-resistant *Staphylococcus aureus* (MRSA) strains used in this study were all initially derived from patients with clinical infections: MRSA 11/11 (USA300), MW2 (USA400), COL (USA100), BMC1001 (USA500), and 300-111 (CC8, *spa* type 4, Iberian clone) (53, 61–65). These prototypical strains encompass the range of clonal complex, *agr* and *SCCmec* genotypes that are in current worldwide clinical circulation. In addition, several of these strains have been previously used in experimental studies of virulence, pathogenesis and antimicrobial responsiveness (MW2; COL; and MRSA 11/11) (5, 23, 53, 66). In experimental IE studies, we also included the well-known MSSA strain, ATCC25923, as a control.

MRSA and MSSA strains were stored at −80°C until thawed for use. They were isolated on tryptic soy agar (TSA) and incubated at 37°C in ambient air. Bacteria were grown for most experiments, including AST, overnight in cation-adjusted Mueller-Hinton Broth (**CA-MHB**; Difco) with the addition of 100 mM Tris (hydroxymethyl-aminomethane) to maintain pH at ~7.3 ± 0.1 throughout all AST testing (Fisher Scientific). In parallel studies, CA-MHB-Tris was supplemented with either 25 mM or 44 mM NaHCO_3_; these NaHCO_3_ concentrations represent physiologic bloodstream concentrations and those found in DMEM, respectively. In parallel control assays, AST testing was also performed in the tissue culture medium, Roswell Park Memorial Institute (**RPMI**) 1640 (Fisher Scientific) supplemented with 10% Luria-Bertani (**LB**) broth. All media was supplemented with 2% NaCl when performing assays in which MRSA cells were exposed to oxacillin.

### Minimum inhibitory concentration (MIC) assays

The MICs of cefazolin and oxacillin were determined according to the Clinical and Laboratory Standards Institute **(CLSI)** guidelines by broth microdilution (12, 13). MRSA were grown overnight in specified media and diluted into the same media containing 2-fold serial dilutions of antibiotics. 2% NaCl was added to all media when performing oxacillin MICs. All MIC values are the mode of at least 6 independent determinations.

For MIC determinations done in the presence of absence of salicylic acid, the following media were used: CA-MHB-100 mM Tris ± 25 or 50 μg/mL salicylic acid.

### Time-kill assays

Cells were grown overnight in specified testing media and diluted to 5 x 10^5^ CFU/mL in 200 μL of same media on a 96-well plate (flat bottom; tissue culture-treated). Based on pilot experiments, cells were incubated at 37°C for 3 h to enter log phase, then diluted to 5 x 10^5^ CFU/mL in 200 μL of the same media, with or without antibiotic, in a 96-well flat bottom plate. Plates were then incubated at 37°C for 24 h. Surviving cells were quantified at 0, 2, 4, 8, and 24 h of incubation and data expressed as log_10_ CFU/ml. A bactericidal effect was defined as a ≥ 3 log_10_ CFU/mL decline in counts at 24 h vs the 0 hr count.

### Population analyses profiles (PAP)

The PAP protocol was modified from published guidelines (67–70). In brief, agar plates were prepared with Mueller Hinton Agar (MHA) supplemented with 100 mM Tris, 2% NaCl, and 44 mM NaHCO_3_ (where indicated), at pH ~7.3 ± 0.1, containing two-fold serial dilutions of cefazolin or oxacillin at concentrations ranging from 0.0625 – 1024 μg/mL. Bacterial cells were grown overnight in broth testing medium and diluted to ~1 × 10^9^ CFU/mL in phosphate buffered saline (PBS). Ten-fold serial dilutions were performed in PBS, and 10 μL of each dilution was plated onto the MHA plates. After 48 h incubation at 30°C, plates were enumerated for viable cells at each drug concentration. The area under the PAP curve (AUC) was calculated by linear approximation.

### Synergy of β-lactams with host defense peptides

For these studies, a prototypical host defense peptide, LL-37, was employed. This cathelicidin peptide is commonly found in large amounts in human epithelial cells, as well as in neutrophils (71, 72), and has been documented to play an important role in innate immunity (73–75). Highly purified LL-37 was purchased commercially from Peptides International (Lexington, KY).

MRSA strains were grown overnight in bicarbonate-free DMEM (Gibco) supplemented with 25 mM NaHCO_3_ and diluted into the same medium supplemented with 150 mM NaCl. Diluted cells were incubated for 3 h at 37°C to enter log phase growth. Log phase cells were then diluted to 1 × 10^3^ CFU/mL with LL-37, cefazolin, or oxacillin alone or in combination; this inoculum has been used standardly in our prior killing assays with host defense peptides (76). Final concentrations of antimicrobials were: 2.5 μg/mL LL-37 (MRSA 11/11), 5 μg/mL LL-37 (MW2 and COL), 0.03125 μg/mL cefazolin and oxacillin (MRSA 11/11), 0.0625 μg/mL cefazolin and oxacillin (MW2 and COL). These antibiotic concentrations were determined after extensive pilot studies as representing individual drug levels that did not cause ≥ 50% killing of this starting MRSA inoculum. Surviving cells were quantified after 4 h incubation with antimicrobials at 37°C, and the percent survival calculated at this time-point as:

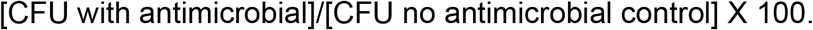

### Isolation of RNA and quantitative real-time PCR (qRT-PCR) analyses

To quantify expression of two key genes involved in the MRSA phenotype *(mecA; sarA),* total RNA was isolated from the study strains following overnight growth in media with and without NaHCO_3_ supplementation using an RNeasy Kit (Qiagen, Valencia, CA) (66). Cells were grown overnight in specified media (CA-MHB 100 mM Tris ± 44 mM NaHCO_3_), then diluted 1:100 into same media and incubated at 37°C overnight. To quantify the combined effect of oxacillin stimulation and NaHCO_3_ exposure on gene expression, cells were grown as specified, but diluted into media containing ½ MIC concentration of oxacillin (with 2% NaCl). qRT-PCR was performed using primers for *mecA, sarA,* and *gyrB* as previously described (66, 77, 78). *gyrB* was used as a housekeeping gene to normalize transcript quantifications. Relative quantification was calculated using the ΔΔC_T_ method. All qRT-PCR gene expression data were determined from two separate biological replicates for each condition, tested in triplicate. Data are presented as the fold-change in gene expression in the presence of NaHCO_3_ exposures when compared to CA-MHB Tris alone for each strain, with CA-MHB Tris gene expression data being normalized to 1.0.

### PBP2a agglutination assays

A semi-quantitative, rapid and reliable latex agglutination method (Seiken, Tokyo, Japan) was used to measure PBP2a production (79), using beads labelled with specific anti-PBP2a antibody (80). Strains were grown overnight in specified media (in the presence or absence of NaHCO_3_ supplementation) at 37°C. Cells were collected by centrifugation, washed once in PBS, resuspended in PBS to an optical density of OD_600nm_ 1.0, and pelleted. Pellets were prepped for PBP2a agglutination as per the manufacturer’s instructions. Agglutination results were scored blindly and separately by two investigators (SCE; LL), and read as high (+++), moderate (++), low (+), or negative (-) based on the presence or absence of an overt agglutination pattern. S. *aureus* ATCC 43300 (MRSA; PBP2a-positive) and ATCC 25923 (MSSA; PBP2a-negative) were used as positive and negative controls, respectively, in all assays.

### Lipase assays

Lipase production is normally repressed by *sarA* (33, 81). As a phenotypic readout for NaHCO_3_-mediated *sarA* repression, a spectrophotometric assay measuring the rate of clearance of a tributyrin emulsion was used to measure lipase activity (82). Strains were grown overnight in specified media (with or without NaHCO_3_ supplementation) at 37°C, then diluted to ~1 x 10^9^ CFU/mL in same media, and filter-sterilized. A 0.5% v/v tributyrin solution (Sigma) was prepared in 100 mM Tris (pH 8.0) + 25 mM CaCl_2_ and emulsified by sonication for 3 min. The tributyrin emulsion was diluted 1:1 with a 0.8% w/v low-gelling-temperature agarose (Sigma), and the suspension maintained at 50°C. Then, 1 mL of the tributyrin suspension was added to 100 μL of each supernatant in a spectrophotometric cuvette, and the OD_450nm_ optical density measured at time 0, 1, 2, 3, 4, and 5 min. Lipase activity was calculated as the rate of clearance normalized to the cell density of each sample. A faster rate of clearance corresponds to greater lipase activity. Data are presented as the fold-change in lipase activity in the presence of NaHCO_3_ as compared to CA-MHB Tris alone for each strain, with CA-MHB Tris-alone lipase activity being normalized to 1.0. As stated above, increases in lipase activity correlate with repression of *sarA* gene ‘tone’ (activity) (33).

### Rabbit model of MRSA infective endocarditis

To verify the *in vivo* translatability of the relationship between NaHCO_3_-responsiveness or NaHCO_3_-nonresponsiveness observed *in vitro*, a well-characterized rabbit model of indwelling catheter-induced aortic valve infective endocarditis (**IE**) was used (66). Rabbits were infected iv at 48 h after catheter placement with 2 x 10^5^ CFU/animal of the indicated strain; this inoculum represents the ID95 for inducing IE as established by extensive pilot experiments for each strain. At 24 h post-infection, animals were randomized into either an untreated control group (sacrificed at this time-point as a therapeutic baseline) or β-lactam-treated groups (100 mg/kg cefazolin or oxacillin, administered by intramuscular injection, t.i.d. for 4 days). These β-lactam treatment strategies encompass: i) dose-regimens used in prior studies of experimental IE (5); and ii) doses that mimic human-like pharmacokinetics in experimental IE (29).

To provide a perspective on the extent of β-lactam-mediated killing *in vivo* in experimental IE among NaHCO_3_-responsive vs. NaHCO_3_-nonresponsive MRSA, we performed a parallel study using the highly cefazolin-susceptible MSSA strain ATCC25923 treated with the same cefazolin treatment regimen employed above for MRSA IE.

In all studies, at 24 h after the last antibiotic treatment, animals were sacrificed, and their cardiac vegetations, kidney, and spleen were removed and quantitatively cultured on TSA. MRSA counts were expressed as mean log_10_ CFU per gram of tissue (± SD). To assess the potential emergence of high-level resistance to either cefazolin or oxacillin for strains 11/11 and MW2 during such β-lactam treatments, the three target tissues were parallel-plated on the above media, but containing 64 μg/ml of the antibiotic-of-interest. The limit of detection in target organ cultures in this model, based on average target tissue weights, is ≤ 2 log_10_ CFU/g.

To determine whether pre-incubation of NaHCO_3_-responsive MRSA in media containing NaHCO_3_ itself might influence the initial induction and/or early progression phases of experimental IE, MRSA 11/11 was grown overnight in CA-MHB 100 mM Tris with either 25 mM or 44 mM NaHCO_3_. Rabbits were then infected iv at 48 h after catheter placement with 2 x 10^5^ CFU/animal with NaHCO_3_-preexposed cells. At 24 h post-infection, rabbits were sacrificed and the same target tissues as above were removed and quantitatively cultured on TSA.

### Statistics

All statistical comparisons were made using a Kruskal-Wallis Single Factor ANOVA test and Pairwise Mann-Whitney *U* test post hoc comparison. Data are presented, unless otherwise indicated, as the sample means ± SD. *P* values < 0.05 were considered statistically significant.

### Study Approval

Female, New Zealand White rabbits, weighing 2.2 – 2.5 kg were used in all animal studies (Irish Farm). Rabbits were maintained in accordance with the American Association for Accreditation of Laboratory Animal Care criteria. The Institutional Animal Care and Use Committee of the Los Angeles Biomedical Research Institute at Harbor–UCLA Medical Center approved all animal study protocols.

## SUPPLEMENTAL METHODS

### Serum antibiotic concentration analyses

Rabbit serum concentrations of cefazolin and oxacillin after a 100 mg/kg i.m. dose were measured using a radial diffusion assay (83). Rabbit blood was collected intravenously 1 h and 2 h post drug administration, and serum was collected via centrifugation. *Bacillus subtilis* strain ATCC6633 was grown overnight in Brain Heart Infusion (BHI) medium and washed twice with PBS. Cells were resuspended in PBS, briefly sonicated, and adjusted to an optical density of OD_600nm_ 0.5 (~1 x 10^8^ CFU/mL). Cells were diluted to a final concentration of 5 x 10^5^ CFU/mL in MHA and poured into petri plates. A radial diffusion punch was used to make wells in the agar plates. Rabbit serum was diluted in a 2-fold dilution series in water, and 60 μL of each dilution was added to the agar wells. A standard plate was also prepared with cefazolin or oxacillin concentrations ranging from 0.0625 - 256 μg/mL in 60 μL water. The plates were incubated at 37 °C for 3 h, after which the serum and drug standards were aspirated out of the wells and 10 mL of TSA was poured over the tops of the plates. Plates were incubated at 37 °C overnight, after which the zones of clearance around the wells were measured. A standard curve was constructed and used to calculate rabbit serum concentrations of cefazolin and oxacillin.

### Blood [HCO_3_^-^] analyses

The CG4+ i-STAT cartridge system (Abbott Point of Care Inc., New Jersey) was used as per manufacturer’s instructions to measure [HCO_3_^-^] concentrations in whole rabbit blood collected from the lateral ear vein. Blood samples were taken from animals at baseline pre-infection, at 24 h post-induction of IE for untreated animals, or after 2 days of β-lactam treatment of animals with IE. Strains MRSA 11/11, MW2, COL, and BMC1001 were used for these studies.

## Author’s contributions

All authors have read and approved the final manuscript. Specific contributions were as follows:

Concept and design: SCE; ASB; YQX.

Collection of data: SCE; WA; LL.

Analysis of data: SCE; ASB; LL; YQX; HFC.

Drafting the manuscript: SCE; ASB; YQX.

Critical revision of the manuscript: SCE; ASB; YQX; LL; WA; HFC.

Final acceptance of the manuscript: All Authors.

Agreed to be accountable for all aspects of the work: All Authors.

## ACKNOWLEDGMENTS

This project was supported in part by the following grants: NIH (NIAID) R01 AI100291 (to HFC); R01 AI039108 (to ASB); R01AI041513; 1R42AI36065-01A1; and R42AI118133 (to YQX).

## Notes

The authors have declared no conflict of interest exists.

